## Supplemental Materials for "Lack of *At*MC1 catalytic activity triggers autoimmunity dependent on NLR stability"

#### APPENDIX FIGURES

**Figure S1. The autoimmune phenotype of *atmc1* mutant plants is specific to *At*MC1 and does not occur in other Type Is or a Type II metacaspase mutants.** Representative images of 40-day-old Wt, atmc1, atmc2 (T-DNA mutant), atmc3-CR#13.3 (CRISPR mutant) and atmc4 (T-DNA mutant) plants grown under short day conditions. Scale bar = 5.5 cm.

**Figure S2. The observed autoimmune phenotype occurs both by overexpression or by expressing $AtMC1^{C220A}$*-GFP* under its native promoter.**

**(A)** Representative images of 40-day-old *atmc1 AtMC1-GFP* and *atmc1* $AtMC1^{C220A}$ *- GFP* plants grown under short day conditions. 2 and 5 independent transgenics in homozygosity for the transgene for *atmc1 AtMC1-GFP* and *atmc1* $AtMC1^{C220A}$*-GFP* plants, respectively, are shown. Scale bar = 5.5 cm.

**(B)** Representative images of 40-day-old *atmc1 proAtMC1::AtMC1 -GFP* and *atmc1* $proAtMC1::AtMC1^{C220A}$ *-GFP* grown under short day conditions. 3 independent transgenics in homozygosity are shown for each genotype. Scale bar = 5.5 cm

**Figure S3. The N-terminal prodomain is required for the autoimmune phenotype caused by catalytically inactive *At*MC1.**

**(A)** Representative images of 40-day-old plants with the indicated genotypes grown under short day conditions. 3 and 2 independent homozygous stable transgenics expressing either prodomainless *At*MC1 (*ΔNAtMC1 -GFP* #7.3, #5.15 and 9.11) or prodomainless *At*MC1 catalytically inactive (*ΔNAtMC1* $^{C220A}$*-GFP* #5.1 and #4.2), respectively, under the control of a 35S constitutive promoter in the *atmc1* mutant background are shown. Scale bar = 5.5 cm.

**(B)** Scheme of *At*MC1, *ΔNAt*MC1, and catalytically inactive *ΔNAt*MC1 ($AtMC1^{C220A}$) proteins fused to GFP. The prodomain, p20 and p10 domains are indicated. The catalytic cysteine (C220) is also indicated.

**(C)** Total protein extracts from the plant genotypes shown in A were run on an SDS-PAGE gel and immuno-blotted against the indicated antisera. Comassie Blue Staining (CBS) of the immunoblotted membranes shows protein levels of Rubisco as a loading control.

**(D)** Plant fresh weight of genotypes shown in **A** (n=12). Different letters indicate statistical difference in fresh weight between genotypes (one-way ANOVA followed by post hoc Tukey, $p$ value < 0.05).

**Figure S4. Non-autoprocessable *At*MC1 variants do not display autoimmunity.**

**(A)** Representative images of 40-day-old plants with the indicated genotypes grown under short day conditions. 3 independent homozygous stable transgenics expressing a $Ca^{2+}$

insensitive variant (*At*MC1$^{DED}$-GFP #9.11, #8.15 and #3.1) from a 35S constitutive promoter in the *atmc1* mutant background are shown. Scale bar = 5.5 cm.

**(B)** Total protein extracts from the plants shown in A were run on an SDS-PAGE gel and immuno-blotted against the indicated antisera.

**(C)** Plant fresh weight of genotypes shown in A (n=12). Different letters indicate statistical difference in fresh weight between genotypes (one-way ANOVA followed by post hoc Tukey, p value < 0.05).

**(D)** Representative images of 40-day-old plants with the indicated genotypes grown under short day conditions. One stable transgenic overexpressing a non-cleavable *At*MC1 variant (*At*MC1$^{R49A}$-GFP #11.8) in the *atmc1* mutant background is shown. Scale bar = 5.5 cm.

**(E)** Total protein extracts from *atmc1 AtMC1$^{DED}$-GFP* #9.11, #3.1 and #8.15 and *atmc1 AtMC1$^{R49A}$-GFP* #11.8 were run on an SDS-PAGE gel and immuno-blotted against the indicated antisera.

**Figure S5. Autoimmunity caused by catalytically inactive *At*MC1 is partially dependent on PAD4 but not SAG101.**

**(A)** Representative images of 40-day-old plants with the indicated genotypes grown under short day conditions. Scale bar= 5.5 cm.

**(B)** Trypan blue staining of an area belonging to the 6$^{th}$ true leaf of the plants shown in A. Scale bar = 0.5 mm.

**(C)** Total protein extracts from the plants shown in A were run on a SDS-PAGE gel and immuno-blotted against the indicated antisera. Comassie Blue Staining (CBS) of the immunoblotted membranes shows protein levels of Rubisco as a loading control.

**(D)** Plant fresh weight of genotypes shown in A (n=12). Different letters indicate statistical difference in fresh weight between genotypes (one-way ANOVA followed by post hoc Tukey, p value < 0.05). Quantification of fresh weight from Wt*, pad4-1* and *sag101-1* were excluded from the fresh weight graph to better appreciate statistical differences between genotypes of interest.

**Figure S6. GO terms representing biological processes derived from significantly enriched peptides that co-immunoprecipitated with *At*MC1$^{C220A}$-GFP.** The most specific term from each family term provided by PANTHER was plotted along with the corresponding gene number, fold enrichment (FE), and FDR (Bonferroni correction for multiple testing) represented as log10. Only GO terms with an FE above 2 and FDR below 0.05 were plotted.

**Figure S7. Immune components that interact with catalytically inactive *At*MC1 are individually not required for the autoimmune phenotype caused by catalytically inactive *At*MC1.**

**(A)** Representative images of 40-day-old plants with the indicated phenotypes grown under short day conditions. Two independent stable transgenics in the $T_2$ generation expressing either DN-RPS2 (*DN-RPS2* #1 and #2) or DN-SSI4 AT5G41750 (*DN-SSI4 AT5G41750* #1 and #2*)* under the control of a 35S constitutive promoter in the *atmc1 AtMC1$^{C220A}$-GFP* background are shown. Scale bar=5.5 cm.

**(B)** Representative images of 50-day-old plants with the indicated phenotypes grown under short day conditions. The *rps2-201c*, *rlp42-2* and *rbohf* mutant alleles were introgressed into the *atmc1 AtMC1$^{C220A}$-GFP* background by conventional crosses and pictures were taken in the F3 offspring. Scale bar= 5.5 cm.

**(C)** Representative images of 40-day-old plants with the indicated phenotypes grown under short day conditions. The *rlp23-1*, *sobir1-12* and *bak1-4* mutant alleles were introgressed into the *atmc1/AtMC1$^{C220A}$-GFP* background by conventional crosses and pictures were taken in the F4 offspring. Scale bar= 5.5 cm.

**Figure S8. Immunoblots showing accumulation of RPS2-HA and SSI4-mCitrine in various *At*MC1 backgrounds.**

**(A)** Western blots of the represented values shown in Figure 6B. Total protein extracts from the plant genotypes were run on an SDS-PAGE gel and immuno-blotted against the indicated antisera. Coomassie Blue staining (CBS) of the immunoblotted membranes shows protein levels of Rubisco as a loading control.

**(B)** Total protein extracts from the indicated plant genotypes were run on an SDS-PAGE gel and immune-blotted against the indicated antisera. Coomassie Blue staining (CBS) staining shows protein levels of Rubisco as loading control.

**Figure S9. The N-terminal prodomain is required for microsomal and puncta localization in catalytically inactive *At*MC1.**

**(A)** Fractionation assays from plant extracts with the indicated plant genotypes. Total (T), Soluble (S, cytoplasmic proteins) and Microsomal (M, total membranes) fractions were run on an SDS-PAGE gel and immunoblotted against the indicated antisera. Coomassie Blue staining (CBS) of the immunoblotted membranes shows protein levels of Rubisco as a loading control.

**(B)** Representative confocal microscopy images from the leaf epidermis of 40-day-old plants grown under short day conditions with the indicated genotypes. Images represent a Z-stack of 12 images taken every 1 µm. Arrows indicate some of the puncta structures formed when

*At*MC1$^{C220A}$ is expressed in an *atmc1* mutant background that are not present in Wt *At*MC1 expressing plants or prodomainless variants (*atmc1 ΔNAtMC1-GFP #5,15 and atmc1 ΔNAtMC1$^{C220A-}$GFP #5,1).* Scale bar = 10 µm.

### FIGURE S1: ASSOCIATED TO FIGURE 1

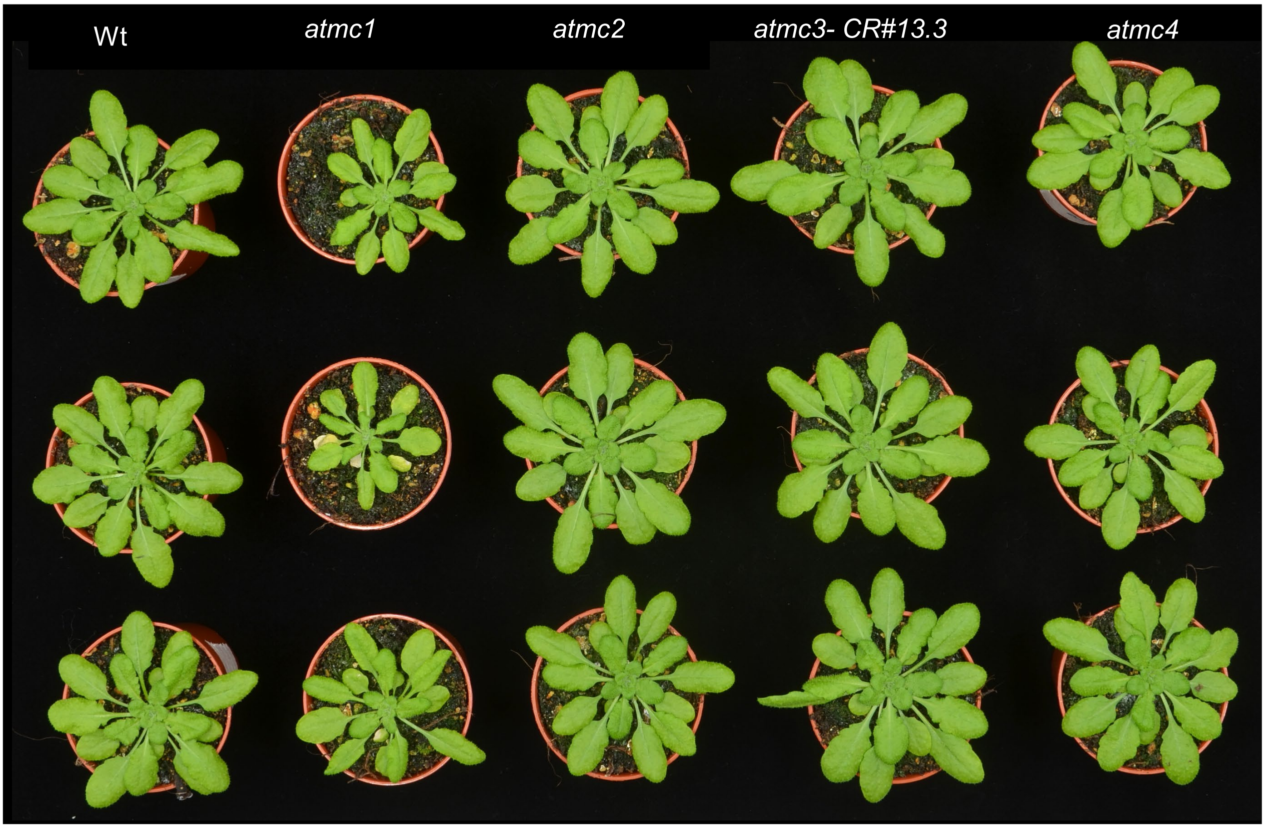

### FIGURE S2: ASSOCIATED TO FIGURE 2

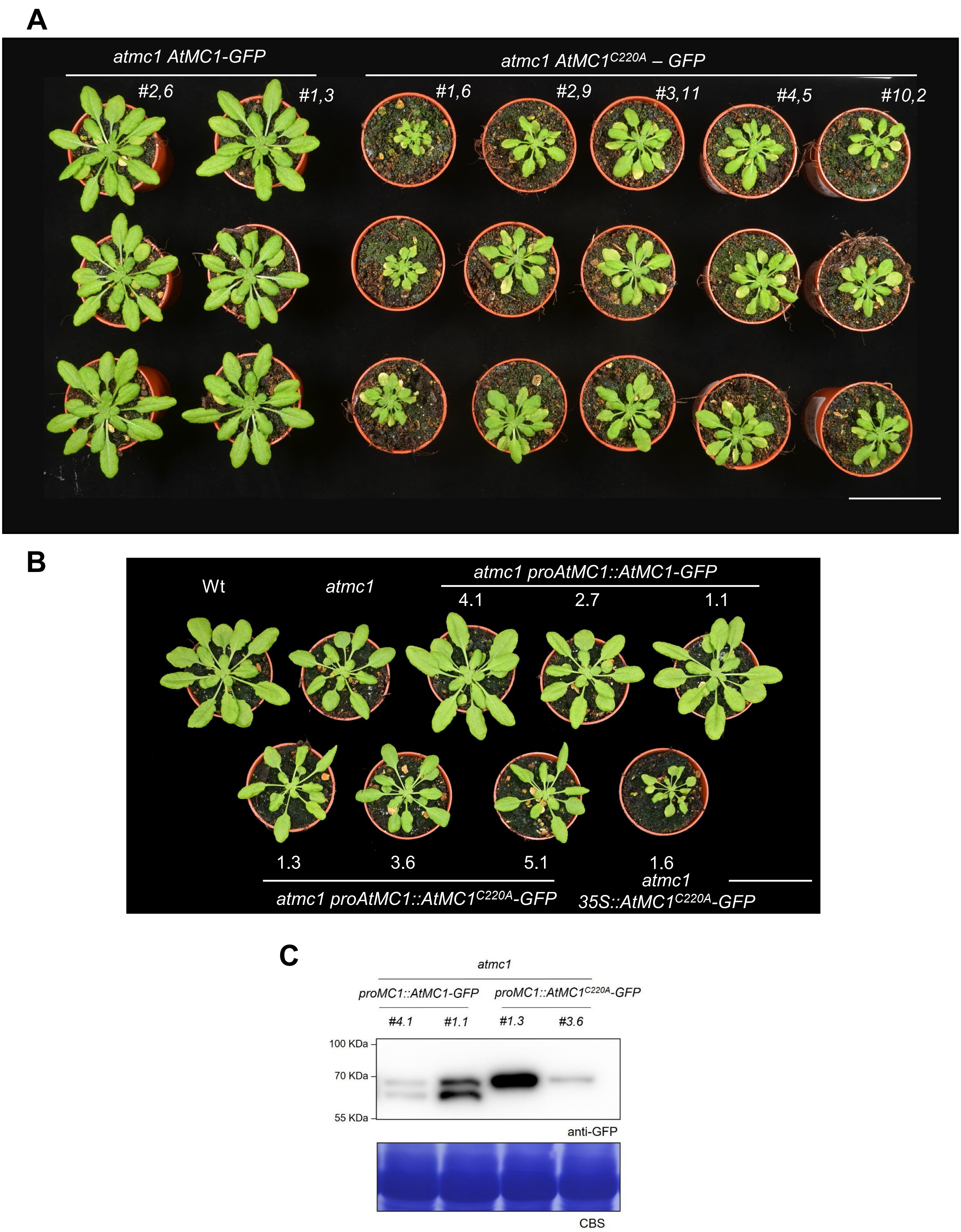

### FIGURE S3: ASSOCIATED TO FIGURE 2

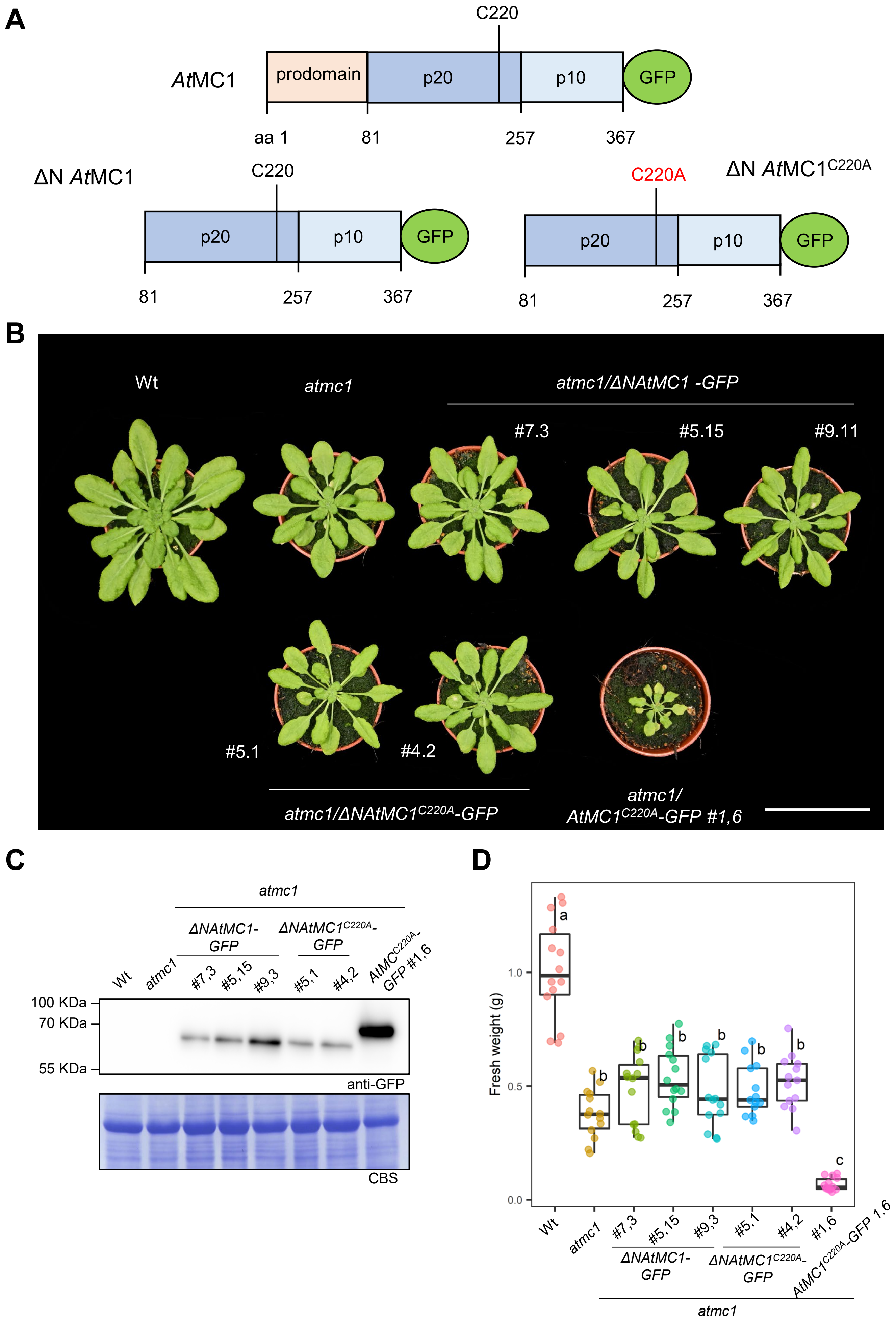

### FIGURE S4: ASSOCIATED TO FIGURE 2

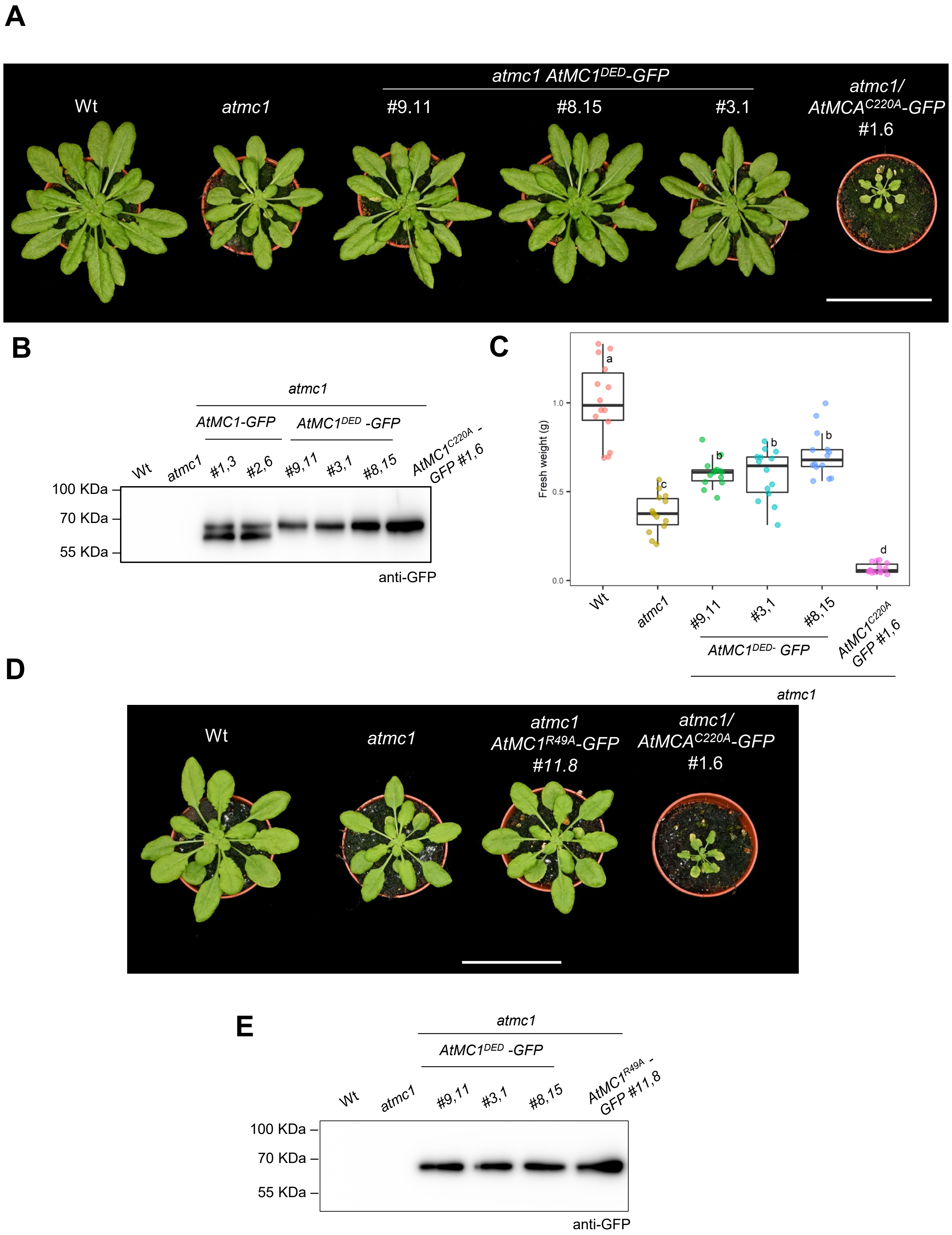

### FIGURE S5: ASSOCIATED TO FIGURE 3

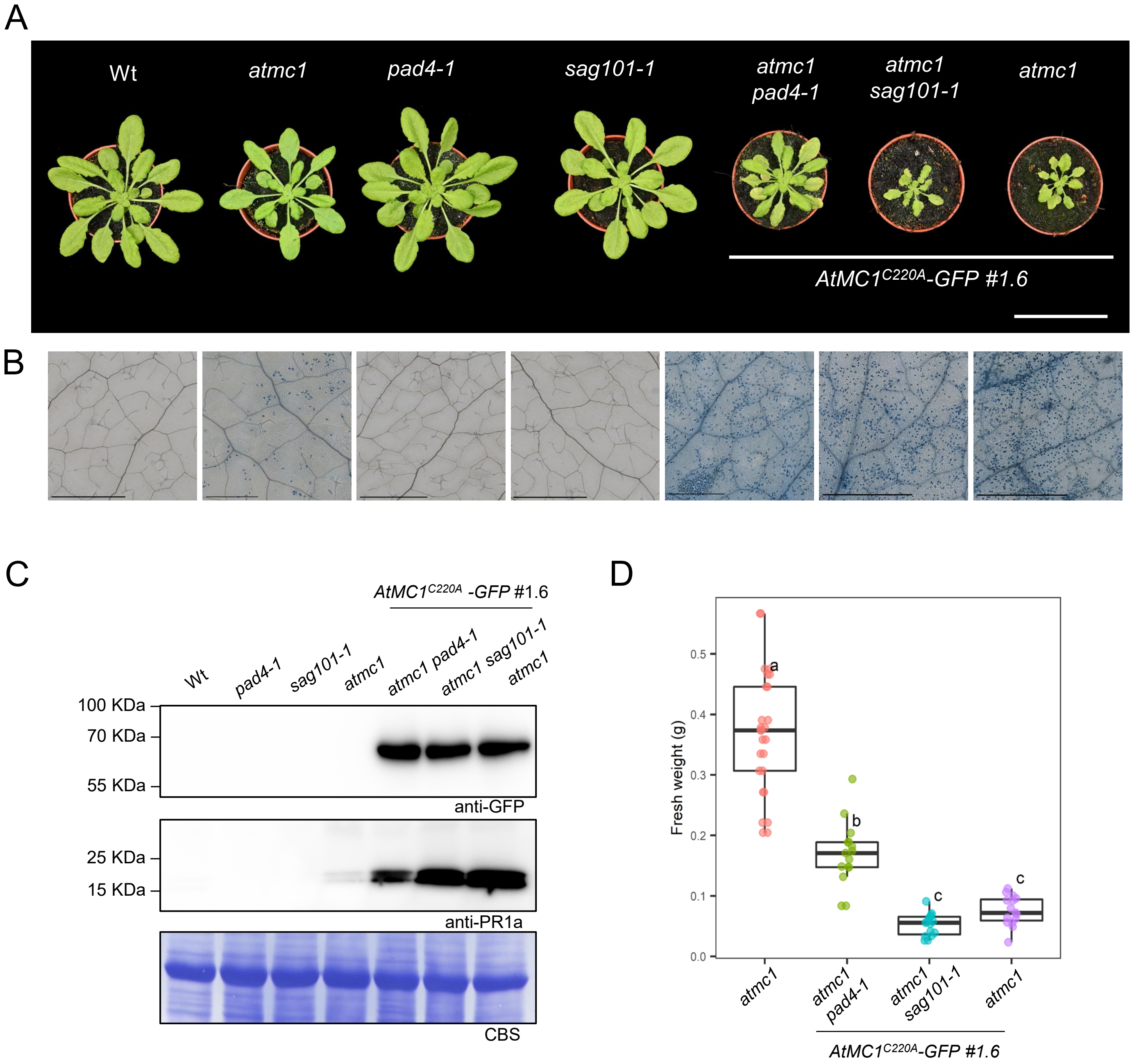

### FIGURE S6: ASSOCIATED TO FIGURE 5

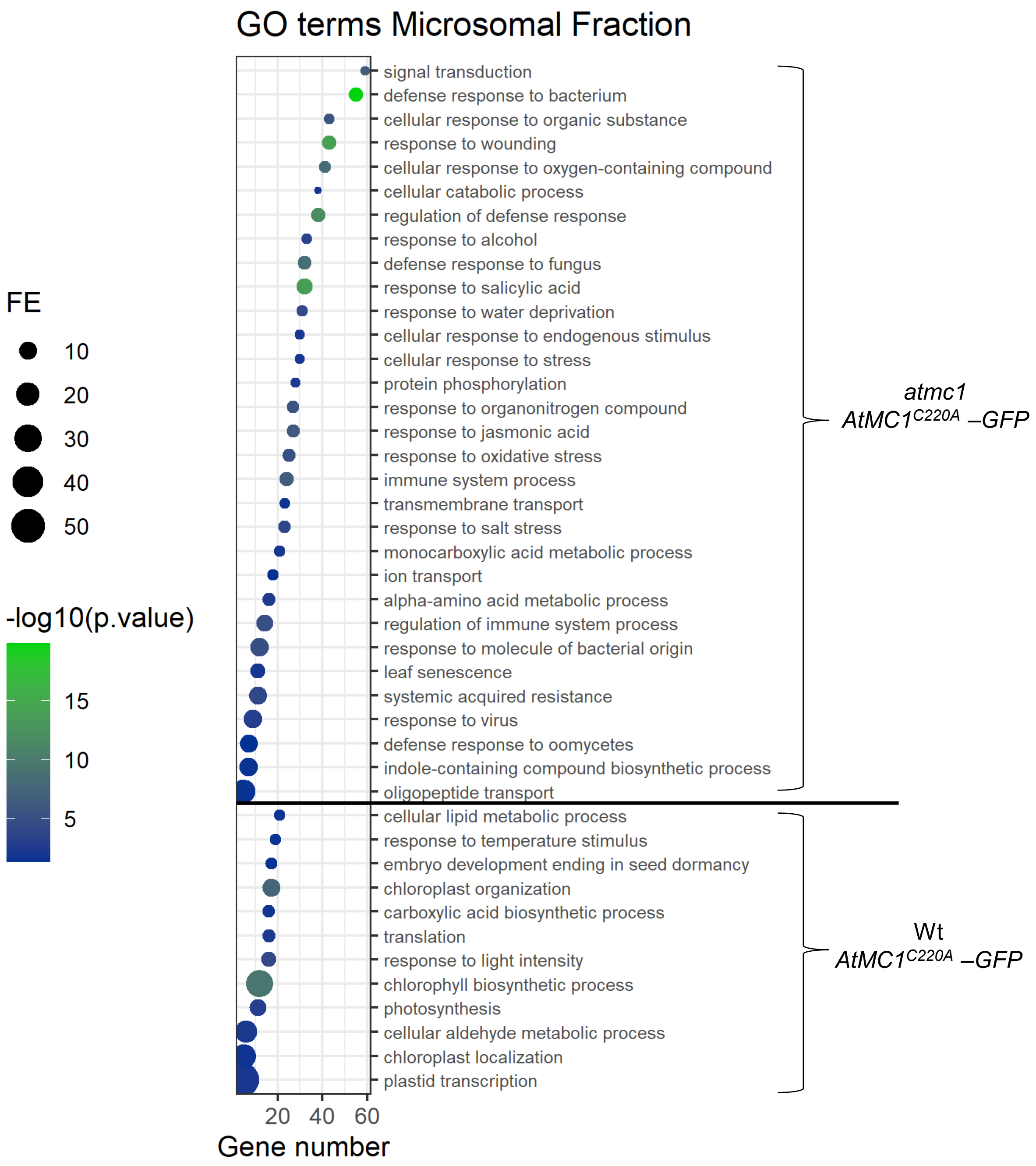

### FIGURE S7: ASSOCIATED TO FIGURE 6

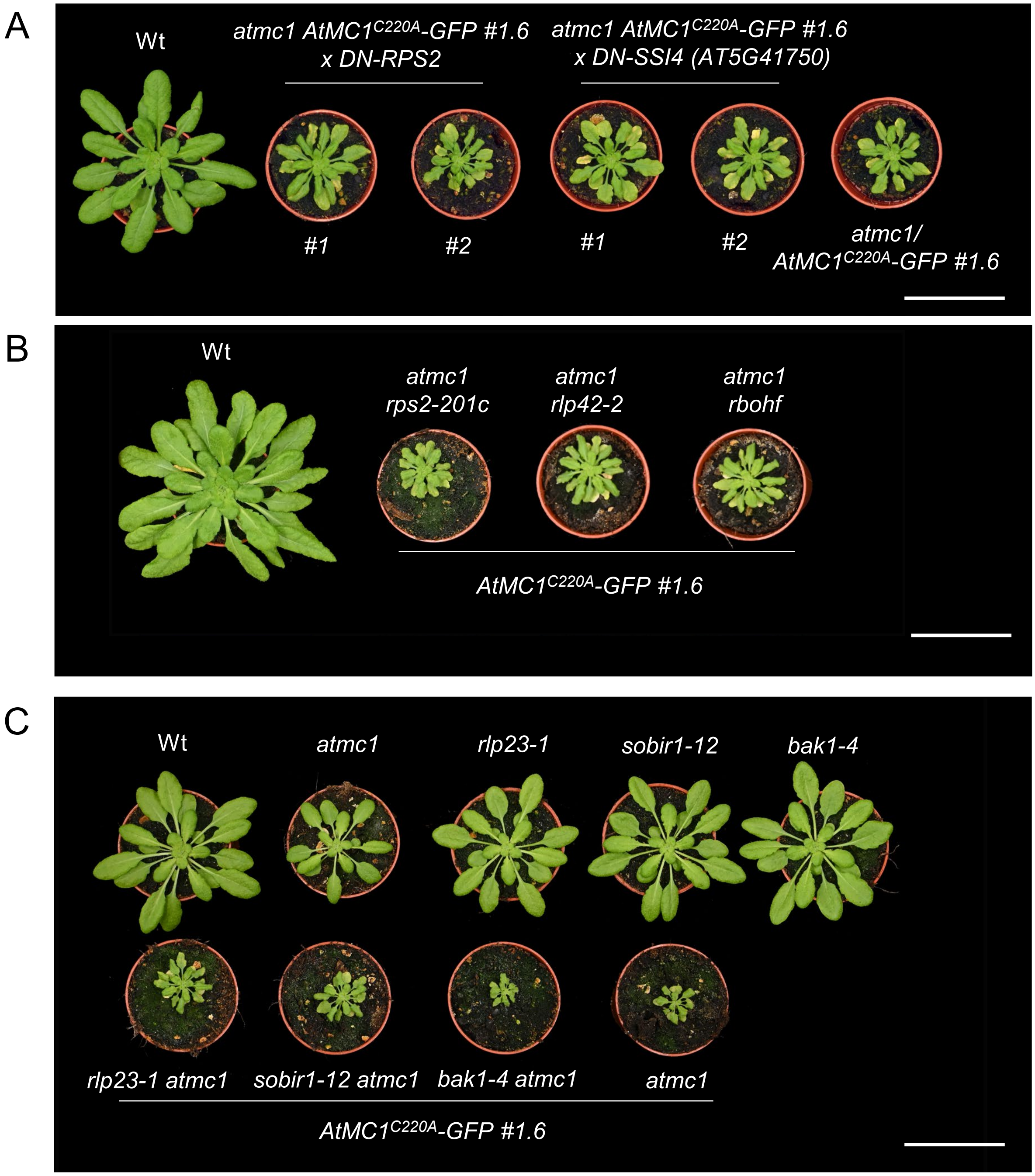

### FIGURE S8: ASSOCIATED TO FIGURE 6

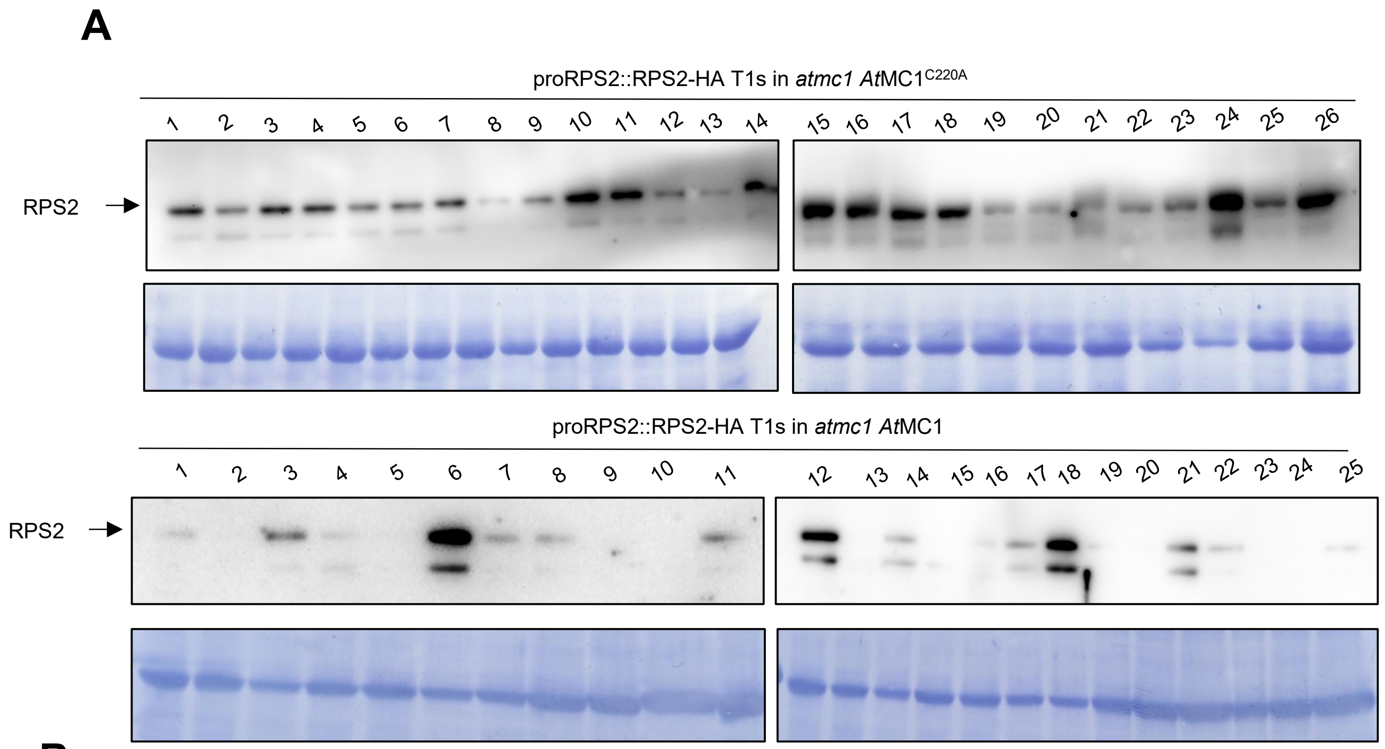


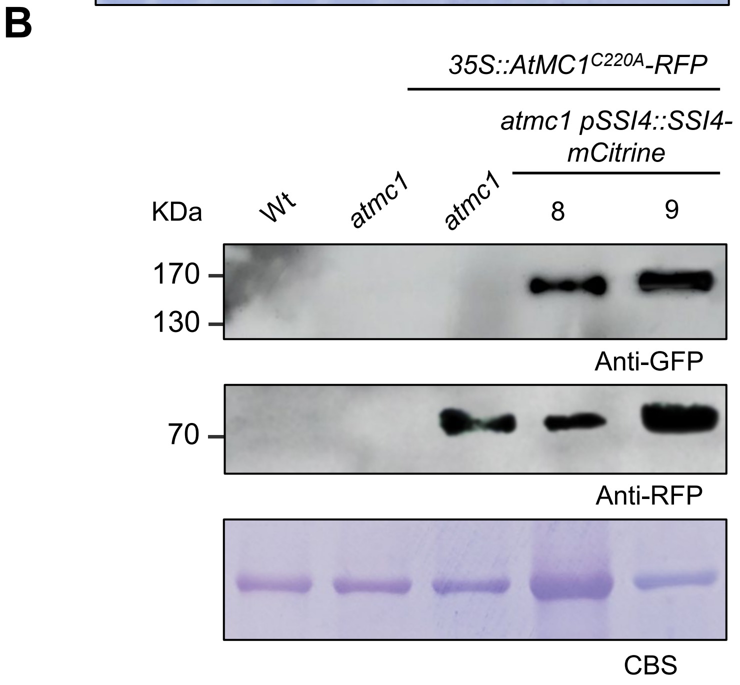

### FIGURE S9: ASSOCIATED TO FIGURE 6

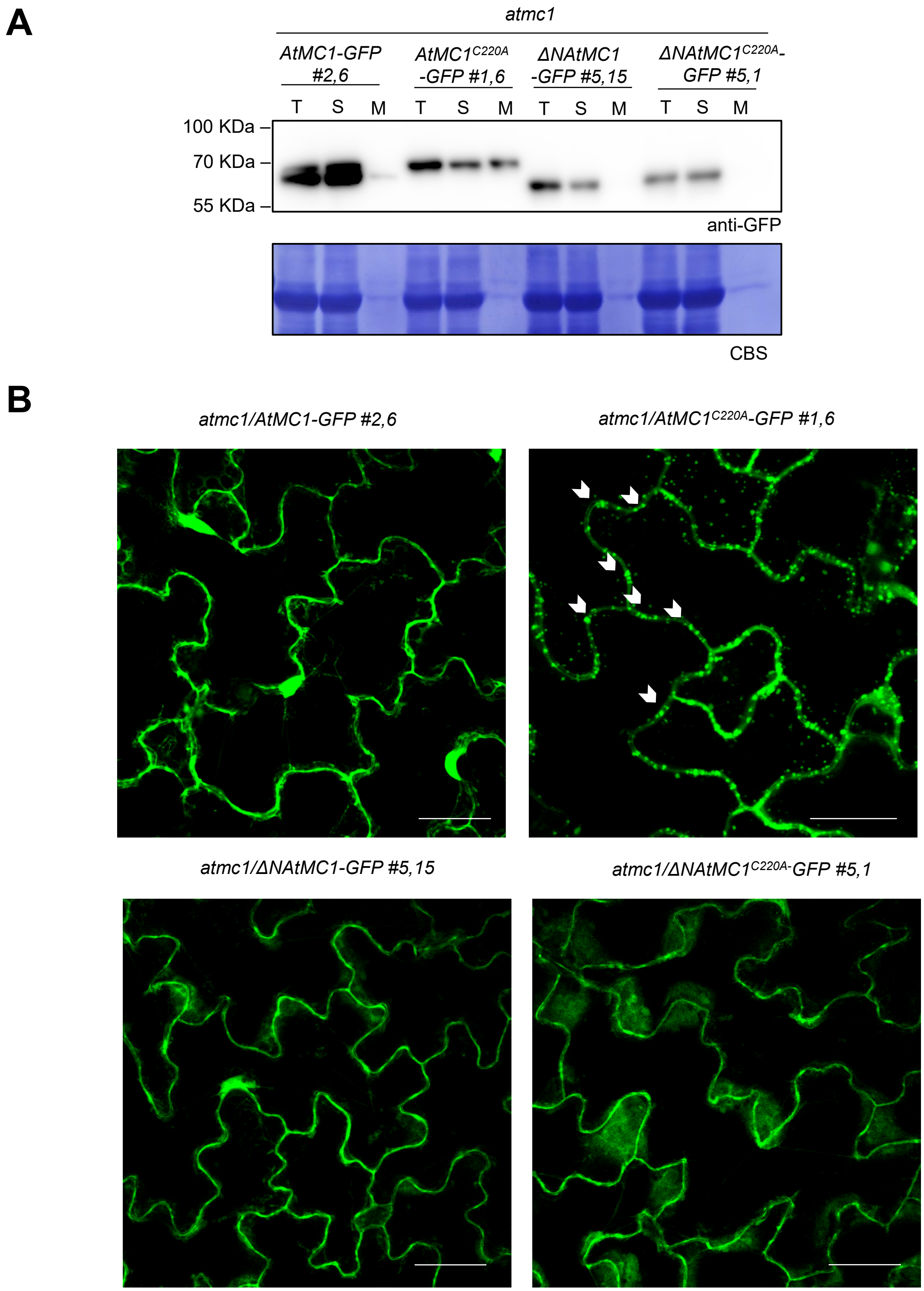

#### APPENDIX TABLES

**Table S1: List of proteins identified by IP-MS/MS as interactors of AtMC1$^{C220A}$, corresponding to the GO term "Defence response to bacterium"**. Significantly enriched peptides that co-immunoprecipitated with *At*MC1$^{C220A}$-GFP in the *atmc1* mutant background. Shown in the table the Log10 pvalue and Log2FC (Bonferroni corrected) provided by PANTHER, as well as the Uniprot ID, gene description and converted gene name.

| Log10 p.value | Log2 FC | Uniprot ID | Gene description | Gene names |
|---|---|---|---|---|
| 3.38 | -4.76 | Q9SMU8 | Peroxidase 34 | PER34 |
| 2.61 | -3.82 | O24658 | Endochitinase At2g43590 | At2g43590 |
| 5.87 | -3.82 | Q6NNH1 | At1g13520 | At1g13520 |
| 3.42 | -3.77 | Q8RXU9 | C2 calcium/ipid-binding plant phosphoribosyltransferase family protein | At4g00700 |
| 2.79 | -3.74 | Q9LMN8 | Wall-associated receptor kinase 3;Wall associated kinase 3 | WAK3 |
| 3.61 | -3.68 | Q9FNL7 | Protein NRT1/PTR FAMILY 5.2 ;Peptide transporter 3 | NPF5.2 |
| 2.75 | -3.22 | O49340 | Cytochrome P45071A12 | CYP71A12 |
| 4.30 | -3.21 | B9DG91 | AT3G13950 protein | At3g13950 |
| 8.54 | -3.20 | O22149 | Probable pectinesterase/pectinestera se inhibitor 17 | PME17 |
| 4.65 | -2.97 | Q8RXN7 | Alpha/beta-Hydro lases su perfamily protein | At2g39420 |
| 3.17 | -2.74 | Q9LY77 | Calcium-transporting ATPase 12, plasma mem brane-type | ACA12 |
| 2.05 | -2.63 | Q9LXJ3 | Uncharacterized protein At3g52710 | F3C22 110 |
| 2.42 | -2.50 | Q8L7G3 | Cysteine-rich receptor-like protein kinase 7 | CRK7 |

| | | | | |
|---|---|---|---|---|
| 2.65 | -2.47 | Q9S9U1 | L-type lectin-domain containing receptor kinase VII.1;Concanavalin A-like lectin protein kinase family protein | LECRK71 |
| 2.15 | -2.45 | Q9FE06 | Protein EXORDIUM-like 2 | EXL2 |
| 3.70 | -2.43 | Q9LFT3 | Endo-1,3(4)-beta-glucanase | F1N13_10 |
| 3.23 | -2.37 | Q9LTM0 | Cytochrome P45071B23 | CYP71B23 |
| 4.21 | -2.32 | P93733 | Phospholipase D beta 1;Phospholipase D | PLDBETA1 |
| 2.68 | -2.30 | F4K2R6 | Calmodu lin-binding protein 60G;Cam-binding protein 60-like G | CBP60G |
| 2.88 | -2.20 | Q9ZV49 | Expressed protein;Transmembrane protein | At2g18690 |
| 2.34 | -2.19 | P24704 | Su peroxide dismutase [Cu-Zn] 1;Superoxide dismutase [Cu-Zn] 3 | CSD1;CSD3 |
| 3.67 | -2.13 | Q1H583 | GDSL esterase/lipase 22 | GLL22 |
| 2.95 | -2.08 | Q8GW35 | At1g58225;Uncharacterized protein | At1g58225 |
| 2.83 | -2.04 | Q8W4G6 | Cysteine-rich receptor-like protein kinase 15 | CRK15 |
| 2.04 | -2.01 | Q9LIB6 | COBRA-like protein 8 | COBL8 |
| 2.31 | -2.00 | Q940G5 | Glucose-6-phosphate 1-epimerase | F14M19.180 |
| 5.02 | -1.94 | Q9C5S9 | Cysteine-rich receptor-like protein kinase 6 | CRK6 |
| 3.27 | -1.92 | Q9ZP16 | Cysteine-rich receptor-like protein kinase 11;Cysteine-rich RLK (RECEPTOR-like protein kinase) 11 | CRK11 |
| 4.53 | -1.92 | Q64477 | G-type lectin S-receptor-likeserine/threonine-protein kinase At2g19130 | |

| | | | | |
|---|---|---|---|---|
| 2.01 | -1.88 | Q9SNC6 | U-box domain-containing protein 13 | PUB13 |
| 3.58 | -1.77 | Q64483 | Senescence-induced receptor-like serine/threon line-protein kinase | SIRK |
| 2.85 | -1.75 | Q8RY67 | Wall-associated receptor kinase-like 14 | WAKL14 |
| 2.33 | -1.69 | Q9FYC2 | Pheophorbide a oxygenase, chloroplasti ic | PAO |
| 2.45 | -1.67 | Q80874 | Internal alternative NAD(P)H-ubiquinone oxidoreductase A2, chondrial | NDA2 |
| 2.38 | -1.66 | Q9FLVO | Protein DOWNY MILDEW RESISTANCE 6 | K18P6.6 |
| 3.25 | -1.59 | Q9LQU2 | Protein PLANT CADMIUM RESISTANCE 1 | PCR1 |
| 3.50 | -1.58 | Q9FNH6 | NDR1/HIN1-like protein 3 | NHL3 |
| 3.14 | -1.56 | Q9LSX5 | Disease resistance protein IR-NBS-LRR class) family | At5g41750 |
| 2.05 | -1.56 | P43296 | Cysteine protease RD19A | RD19A |
| 2.43 | -1.52 | Q42484 | Disease resistance protein RPS2 | RPS2 |
| 4.56 | -1.48 | Q9CAR7 | Hypersensitive-induced response protein 2 | HIR2 |
| 3.86 | -1.46 | Q48837 | Receptor like protein kinase S.2 | LECRKS2 |
| 2.42 | -1.37 | O9FKH6 | Cytochrome b561 and DOMON domain-containing protein At5g35735 | |
| 2.08 | -1.35 | A0A178WLG 7 | Leucine-rich repeat protein kin ase family protein;Leucine-rich repeat protein kinase family protein | At1g51790 |

| | | | | |
|---|---|---|---|---|
| 2.42 | -1.28 | Q9SY59 | NF-X1-type zinc finger protein NFXL1;NF-X-like 1;NF-X-like 1 | NFXL1 |
| 2.55 | -1.26 | O80517 | Uclacyanin-2 | |
| 2.16 | -1.18 | Q9SU71 | Protein EDS1B | EDS1B |
| 4.94 | -1.15 | Q8H199 | Cysteine-rich receptor-like protein kinase 14 | CRK14 |
| 2.21 | -1.14 | F4KA19 | Calcium-binding EF hand family protein | At5g28830 |
| 2.41 | -1.13 | Q93Y40 | Oxysterol-binding protein-related protein 3C | ORP3C |
| 3.60 | -1.12 | Q9LZU4 | Cysteine-rich receptor-like protein kinase 4 | CRK4 |
| 2.96 | -1.02 | Q37145 | Calcium-transporting ATPase 1;Calcium-transporting ATPase;Calcium-transporting ATPase | ACA1 |
| 3.69 | -1.01 | Q8S8Q6 | Tetraspanin-8 | TET8 |

**Table S2**: DN-NLRs carrying P-loop mutations transformed in the autoimmune background *atmc1 AtMC1*$^{C220A}$*-GFP*. In green, TNLs and blue CNLs. The battery of DN negative NLRs was produced by Lolle and co-workers (Lolle et al., 2017). No rescues in $T_1$ were achieved after screening for suppression of the phenotype.

| Accession number (TNLs) | Accession number (CNLs) |
|---|---|
| At1g17600 | At1g12210 |
| At1g17610 | At1g12220 |
| At1g27170 | At1g12280 |
| At1g31540 | At1g12290 |
| At1g56520 | At1g52660 |
| At1g56540 | At5g63020 |
| At1g63730 | At4g33300 |
| At1g69740 | At1g53350 |
| At1g63750 | At1g63360 |
| At1g63870 | At1g33560 |
| At1g64070 | At5g43830 |
| At1g65850 | At5g43740 |
| At1g66090 | At3g14460 |
| At1g69550 | At3g14470 |
| At1g72840 | At4g14610 |
| At1g72850 | At5g04720 |
| At1g72860 | At5g05400 |
| At1g72870 | At5g35450 |
| At1g72900 | At5g45510 |
| At1g72910 | At1g15890 |
| At1g72940 | At3g46530 |
| At1g72950 | At5g66630 |
| At2g16870 | At5g66900 |
| At2g17050 | At5g66910 |
| At3g04210 | At3g07040 |
| At3g04220 | At4g27190 |
| At3g44400 | At4g27220 |

| | |
|---|---|
| At3g44480 | At1g58390 |
| At3g44630 | At1g58410 |
| At3g44670 | At1g58807 |
| At3g51560 | At1g58848 |
| At3g51570 | At1g59124 |
| At4g09360 | At1g59218 |
| At4g09420 | At1g59620 |
| At4g12010 | At1g17615 |
| At4g16940 | At1g61190 |
| At4g16950 | At1g63350 |
| At4g16960 | At1g63880 |
| At4g19500 | At3g15700 |
| At4g19510 | At3g46710 |
| At4g19530 | At3g46730 |
| At4g23440 | At4g14370 |
| At5g11250 | At4g19050 |
| At5g17680 | At4g26090 |
| At5g17880 | At5g47250 |
| At5g17970 | At5g47260 |
| At5g18350 | At5g47280 |
| At5g18360 | At5g56220 |
| At5g18370 | |
| At5g22690 | |
| At5g36930 | |
| At5g38340 | |
| At5g38350 | |
| At5g38850 | |
| At5g40060 | |
| At5g40090 | |
| At5g40100 | |
| At5g40910 | |
| At5g40920 | |
| At5g41540 | |
| At5g41550 | |
| At5g41740 | |
| At5g41750 | |

| |
|---|
| At5g44510 |
| At5g45050 |
| At5g45060 |
| At5g45200 |
| At5g45230 |
| At5g45240 |
| At5g45260 |
| At5g46260 |
| At5g46270 |
| At5g46450 |
| At5g46470 |
| At5g46510 |
| At5g46520 |
| At5g48770 |
| At5g48780 |
| At5g49140 |
| At5g51630 |
| At5g58120 |
| At2g14080 |
| At4g36150 |
| At4g16890 |
| RPS4 |
| At1g50180 |
| At4g10780 |
| at1g10920 |
| At3g50950 |
| At1g61180 |
| At1g61310 |

**Table S3. Stress granule markers tested in this study.** List of constructs corresponding to stress granule marker genes that were attempted to be transformed into the described backgrounds.

| **Construct** | **Genetic background** | **Reference** |
|---|---|---|
| *35S::RBP47-RFP* | *atmc1 35S::AtMC1$^{C220A}$-GFP* | (Weber *et al*, 2008) |
| *35S::PAB1-RFP* | *atmc1 35S::AtMC1$^{C220A}$-GFP* | (Maruri-López *et al*, 2021) |
| *35S::TSN2-RFP* | *atmc1 35S::AtMC1$^{C220A}$-GFP* | (Gutierrez-Beltran *et al*, 2021) |
| *35S::eI4FE-RFP* | *atmc1 35S::AtMC1$^{C220A}$-GFP* | (Weber *et al*, 2008) |
| *35S::PAB1-RFP* | *atmc1 35S::AtMC1-GFP* | (Maruri-López *et al*, 2021) |
| *35S::TSN2- RFP* | *atmc1 35S::AtMC1-GFP* | (Gutierrez-Beltran *et al*, 2021) |

**Table S4: List of Arabidopsis lines used in this study.**

| Arabidopsis seeds | Accession number | Source or reference |
|---|---|---|
| *atmc1* (GABI-Kat: GK-096A10) | AT1G02170 | (Coll *et al*, 2010) |
| *atmc1-CR#1* | AT1G02170 | This study |
| *atmc2* (SALK_009045) | AT4G25110 | (Coll *et al*, 2010) |
| *atmc3-CR#13.3* | AT5G64240 | (Pitsili *et al*, 2022) |
| *atmc4* (SAIL_856_D0) | AT1G79340 | (Watanabe & Lam, 2011) |
| *eds1-12* | AT3G48090 | (Ordon *et al*, 2017) |
| *sid2-1* | AT1G74710 | (Wildermuth *et al*, 2001) |
| *pad4-1* | AT3G52430 | (Jirage *et al*, 1999) |
| *sag101-1* | AT5G14930 | (Feys *et al*, 2005) |
| *nrg1.1 nrg1.2* | AT5G66900, AT5G66910 | (Castel *et al*, 2019, 201) |
| *helperless (nrg1.1 nrg1.2, adr1, adr1-L1, adr1-L2)* | AT5G66900, AT5G66910 AT1G33560, AT4G33300, AT5G04720 | (Saile *et al*, 2020) |
| *rps2-201-c* | AT4G26090 | (Kunkel *et al*, 1993) |
| *rlp42-2* | AT3G25020 | (Wang *et al*, 2008) |
| *rbohf* | AT1G64060 | (Torres *et al*, 2002) |
| *bak1-4* | AT4G33430 | (Dressano *et al*, 2017) |

| | | |
|---|---|---|
| *rlp23-1* | AT2G32680 | (Wang *et al*, 2008) |
| *sobir1-12* | AT2G31880 | (Gao *et al*, 2009) |
| *atg2-1 (SALK_076727)* | AT3G19190 | (Thompson *et al*, 2005) |
| *atg5-1 (SAIL_129B07)* | AT5G17290 | (Yoshimoto *et al*, 2009) |
| *atmc1 eds1-12* | AT1G02170, AT3G48090 | This study |
| *atmc1 sid2-1* | AT1G02170, AT1G74710 | This study |
| *atmc1 pad4-1* | AT1G02170, AT3G52430 | This study |
| *atmc1 nrg1.1 nrg1.2* | AT1G02170, AT5G66900, AT5G66910 | This study |
| *atmc1 35S::AtMC1-GFP* | AT1G02170 | This study |
| *atmc1 35S::AtMC1$^{C220A}$-GFP* | AT1G02170 | This study |
| *atmc1 proMC1::AtMC1-GFP* | AT1G02170 | This study |
| *atmc1 proMC1::AtMC1$^{C220A}$-GFP* | AT1G02170 | This study |
| Wt*; 35S::AtMC1$^{C220A}$-GFP* | AT1G02170 | This study |
| *atmc1-CR#2 35S::AtMC1$^{C220A}$-GFP* | AT1G02170 | This study |
| *atmc1 35S::ΔNAtMC1 -GFP* | AT1G02170 | This study |
| *atmc1 35S::ΔNAtMC1 $^{C220A}$ -GFP* | AT1G02170 | This study |
| *atmc1 35S::AtMC1$^{DEED}$-GFP* | AT1G02170 | This study |
| *atmc1 35S::AtMC1$^{R49A}$-GFP* | AT1G02170 | This study |
| *atmc1 eds1-12 35S::AtMC1$^{C220A}$-GFP* | AT1G02170, AT3G48090 | This study |
| *atmc1 sid2-1 35S::AtMC1$^{C220A}$-GFP* | AT1G02170, AT1G74710 | This study |
| *atmc1 sag101-1 35S::AtMC1$^{C220A}$-GFP* | AT1G02170, AT5G14930 | This study |
| *atmc1 pad4-1 35S::AtMC1$^{C220A}$-GFP* | AT1G02170, AT3G52430 | This study |

| | | |
|---|---|---|
| *atmc1 nrg1.1 nrg1.2 35S::AtMC1*$^{C220A}$*-GFP* | AT1G02170, AT5G66900, AT5G66910 | This study |
| *atmc1-CR#2 helperless 35S::AtMC1*$^{C220A}$*-GFP* | AT1G02170, AT5G66900, AT5G66910 AT1G33560, AT4G33300, AT5G04720 | This study |
| *atmc1 atg2-1 35S::AtMC1*$^{C220A}$*-GFP* | AT1G02170, AT3G19190 | This study |
| *atmc1 atg5-1 35S::AtMC1*$^{C220A}$*-GFP* | AT1G02170, AT5G17290 | This study |
| *atmc1 35S::HA-AtSNIPER x 35S::AtMC1*$^{C220A}$*-GFP* | AT1G02170, AT1G14200 | This study |
| *atmc1 UBQ-mCherry-ATG8a x 35S::AtMC1*$^{C220A}$*-GFP* | AT1G02170, AT4G21980 | This study |
| *atmc1 UBQ- mCherry-AtRabA5d x 35S::AtMC1*$^{C220A}$*-GFP* | AT1G02170, AT2G31680 | This study |
| *atmc1 UBQ-mCherry-AtRabG3C x 35S::AtMC1*$^{C220A}$*-GFP* | AT1G02170, AT3G16100 | This study |
| *atmc2 35S::AtMC2-GFP* | AT4G25110 | This study |
| *atmc2 35S::AtMC2*$^{C256A}$*-GFP* | AT4G25110 | This study |
| *atmc1 35S::DN-AtRPS2 x 35S::AtMC1*$^{C220A}$*-GFP* | AT4G26090, AT1G02170 | This study |
| *atmc1 35S::DN-SSI4 (AT5G41750) x 35S::AtMC1*$^{C220A}$*-GFP* | AT5G41750, AT1G02170 | This study |
| *atmc1 rps2-201c 35S::AtMC1*$^{C220A}$*-GFP* | AT1G02170, AT4G26090 | This study |
| *atmc1 rlp42-2 35S::AtMC1*$^{C220A}$*-GFP* | AT1G02170, AT3G25020 | This study |
| *atmc1 rbohf 35S::AtMC1*$^{C220A}$*-GFP* | AT1G02170, AT1G64060 | This study |
| *atmc1 rlp23-1 35S::AtMC1*$^{C220A}$*-GFP* | AT1G02170, AT2G32680 | This study |
| *atmc1 bak1-4 35S::AtMC1*$^{C220A}$*-GFP* | AT1G02170, | This study |

| | | |
|---|---|---|
| | AT4G33430 | |
| *atmc1 sobir1-12 35S::AtMC1$^{C220A}$-GFP* | AT1G02170, AT2G31880 | This study |
| *atmc1 35S::AtMC1$^{C220A}$-RFP* | AT1G02170 | This study |
| *atmc1 35S::AtMC1$^{C220A}$-RFP proSSI4::SSI4-mCitrine (AT5G41750)* | AT1G02170, AT5G41750 | This study |
| *atmc1 35S::AtMC1C220A-GFP proRPS2:RPS2-HA* | AT1G02170, AT4G26090 | This study |
| *atmc1 35S::AtMC1-GFP proRPS2:RPS2-HA* | AT1G02170, AT4G26090 | This study |

**Table S5: List of Plasmids used in this study.**

| Name | Accesion number | Backbone | Source of Reference | Additional information |
|---|---|---|---|---|
| *35S::AtMC1-GFP* | AT1G02170 | pZ003 | This study | Hygromycin Resistance |
| *35S::AtMC1$^{C220A}$-GFP* | AT1G02170 | pZ003 | This study | Hygromycin Resistance |
| *proMC1::AtMC1-GFP* | AT1G02170 | pZ003 | This study | Hygromycin Resistance |
| *proMC1::AtMC1$^{C220A}$-GFP* | AT1G02170 | pZ003 | This study | Hygromycin Resistance |
| *35S::ΔNAtMC1 -GFP* | AT1G02170 | pZ003 | This study | Hygromycin Resistance |
| *35S::ΔNAtMC1 $^{C220A}$ -GFP* | AT1G02170 | pZ003 | This study | Hygromycin Resistance |
| *35S::AtMC1$^{DEED}$-GFP* | AT1G02170 | pZ003 | This study | Hygromycin Resistance |
| *35S::AtMC1$^{R49A}$-GFP* | AT1G02170 | pZ003 | This study | Hygromycin Resistance |
| *35S::HA-AtSNIPER* | AT1G14200 | pZ003 | This study | Fast Red selection |
| *UBQ- mCherry-AtRabA5d* | AT2G31680 | pZ003 | This study | BASTA Resistance |
| *UBQ-mCherry-AtRabG3C* | AT3G16100 | pZ003 | This study | BASTA Resistance |
| *UBQ-mCherry-ATG8a* | AT4G21980 | pZ003 | Gift from Yasin Dagdas´ lab | Fast Red selection |
| *35S::AtMC2-GFP* | AT4G25110 | pZ003 | This study | Hygromycin Resistance |
| *35S::AtMC2$^{C256A}$-GFP* | AT4G25110 | pZ003 | This study | Hygromycin Resistance |
| *35S::DN-AtRPS2* | AT4G26090 | pUSER007 | (Lolle *et al*, 2017) | BASTA Resistance |

| | | | | |
|---|---|---|---|---|
| *35S::DN-AtSSI4 (AT5G41750)* | AT5G41750 | pUSER007 | (Lolle *et al*, 2017) | BASTA Resistance |
| *35S::GFP* | - | pZ003 | This study | BASTA Resistance |
| *pOCS::AtRPS2-HA* | AT4G26090 | | Gift from Farid El Kasmi´s lab | BASTA Resistance |
| *proRPS2-RPS2-HA* | AT4G26090 | pZ001 | Gift from Farid El Kasmi´s lab | Fast Red selection |
| *35S:: AtSSI4 (AT5G41750)-3xHA* | AT5G41750 | pGWB514 | This study | Hygromycin Resistance |
| *35S::10xcMyc-AtRLP42* | AT3G25020 | pGWB521 | Gift from Thorsten Nürnberger´s lab | Hygromycin Resistance |
| *35S::FLAG-RBOHF* | AT1G64060 | pBin19g | Gift from Cyril Zipfel´s lab | BASTA Resistance |
| *35S::10xcMyc-AtSOBIR1* | AT2G31880 | pGWB521 | Gift from Thorsten Nürnberger´s lab | Hygromycin Resistance |
| *proAtSSI4::AtSSI4 (AT5G41750)-mCitrine* | AT5G41750 | pB7m34GW | This study | BASTA Resistance |
| *proAtSSI4::AtSSI4 (AT5G41750)-mCherry-FLAG* | AT5G41750 | pB7m34GW | This study | BASTA Resistance |

**Table S6: List of primers used in this study.**

| Primer name | Sequence | Purpose |
|---|---|---|
| AtMC1 F3 | GCGTCACCTTCTCATCAACA | Genotyping |
| AtMC1 R3 | ACGGTACCACTATGGCAAGC | Genotyping |
| GABI LB (KIRK) | ATATTGACCATCATACTCATTGC | Genotyping |
| AtMC2 LP | TCCAAACTTCTGCAATGAAGG | Genotyping |
| AtMC2 RP | ATGACACCTGAAGTCCTGTGG | Genotyping |
| LBb1.3 | ATTTTGCCGATTTCGGAAC | Genotyping |
| sid2-1 F | TGTCTGCAGTGAAGCTTTGG | Caps genotyping (MfeI) |
| sid2-1 R | CACAAACAGCTGGAGTTGGA | Caps genotyping (MfeI) |
| EDS1 959 | AACTAGCATACAGAGGGGCA | Genotyping |
| EDS1 960 | GCTGAGAGAAATCGAACCGG | Genotyping |
| EDS1 JG08 | AAAGAAGACAACATTGATCTATATCTATTCTCTTTTCTT | Genotyping |
| PAD4 F | GCGATGCATCAGAAGAG | Caps genotyping BsmF1 |
| PAD4 F | TTAGCCCAAAAGCAAGTATC | Caps genotyping BsmF1 |
| SAG101 MW29 | ATGCAAGGAGGTCAAGATCG | Genotyping |
| SAG101 MW43 | TTGTGACTTACCATAACTCTCG | Genotyping |
| dSpm11 | GGTGCAGCAAAACCCACACTTTTACTTC | Genotyping |
| NRG1 FEK_1070 | GCATCTCCACCTCTTCACA | dCaps Genotyping (AvaII) |
| NRG1 FEK_1071 | CTGAAGAAATGAACCCATGT | dCaps Genotyping (Ava II) |
| *rps2-201-c F* | GAATCTTAGAAAACTGAAGCATCTGG | dCaps Genotyping (RsaI) |
| *rps2-201-c R* | AGTTGTGAAGGCTGTGTAACGTCA | dCaps Genotyping (RsaI) |
| *rlp42-2 LP* | GTCCGAAGGGAAATCTCTTTG | Genotyping |

| | | |
|---|---|---|
| *rlp42-2 RP* | TGGAGTGTTACTTGGATTGGC | Genotyping |
| *rbohf F* MAT 171F | CTTCCGATATCCTTCAACCAACTC | Genotyping |
| *rbohf R* MAT 212F | CGAAGAAGATCTGGAGACGAGA | Genotyping |
| *sobir1-12 LP* | GGAGCCATAGGAGGAACAATC | Genotyping |
| *sobir1-12 RP* | TGACATCTTTACTGTTCGGCC | Genotyping |
| *atg5-1 F* DH417 | ATTCACTTCCTCCTGGTGAAG | Genotyping |
| *atg5-1 R DH418* | TTGTGCCTGCAGGATAAGCG | Genotyping |
| *atg2-1 LP* | GTGGGGCTCATAGCTTAGACC | Genotyping |
| *atg2-1 RP* | TCGAGTGATTCTGTGGTTTCC | Genotyping |
| *AtMC1 pGB000 F* | aacaGGTCTCaaacaATGTACCCGCCACCTCCCTCAAG | Cloning |
| *AtMC1 pGB000 R* | aacaGGTCTCtagccgaGAGTGAAAGGCTTTGCATAGACATCGAATGTTTGG | Cloning |
| *proAtMC1 pGA000 F* | AACAGGTCTCAACCTGCTCGGATATCTGATTCTCCATGT | Cloning |
| *proAtMC1 pGA000 R* | AACAGGTCTCTTGTTTATTATTCTCGGAAGGGAGGGAAT | Cloning |
| *$AtMC1^{C220A}$ F* | CTCCATTCAATTATCGATGCTGCCCATAGTGGTACCGTTCTGG | Cloning (Site-directed Mutagenesis) |
| *$AtMC1^{C220A}$ R* | CCAGAACGGTACCACTATGGGCAGCATCGATAATTGAATGGAG | Cloning (Site-directed Mutagenesis) |
| *ΔNAtMC1 F* | aacaGGTCTCaaacaATGTTCTCTCGCCACGAGCTCAAAGGCTG | Cloning (Site-directed Mutagenesis) |
| *$AtMC1^{DEED}$ F* | GTCAAAGAAACTACAACGGTGCCGCCGTTGCCGGCTATGATGAAACACTCTG | Cloning (Site-directed Mutagenesis) |
| *$AtMC1^{DEED}$ R* | CAGAGTGTTTCATCATAGCCGGCAACGGCGGCACCGTTGTAGTTTCTTTGAC | Cloning (Site-directed Mutagenesis) |

| | | |
|---|---|---|
| *AtMC1$^{R49A}$ F* | TACTCATATCGCCGACCCTGCTACCGCCCCTCCTCCGCA | Cloning (Site-directed Mutagenesis) |
| *AtMC1$^{R49A}$ R* | GTTGCGGAGGAGGGGCGGTAGCAGGGTCGGCGATATGAG | Cloning (Site-directed Mutagenesis) |
| *HA-SNIPER F* | ATGTATCCGTATGATGTTCCGGATTATGCAATGTCTTCTGAGAATGATTTC | Cloning |
| *HA-SNIPER pGB0000 F* | aacaGGTCTCaaacaATGTATCCGTATGATGTTCCGGATTAT | Cloning |
| *AtSNIPER pGB0000 R* | aacaGGTCTCtagccTTAGTTTCTTCTGTCGCCGG | Cloning |
| *AtRabA5d pGC0000 F* | aacaGGTCTCaggctcaacaATGTCGTCCGATGACGAAGGAGGAG | Cloning |
| *AtRabA5d pGC0000 R* | aacaGGTCTCtctgaTCACGAGGAAGAACAGCAAGAGAAAC | Cloning |
| *AtRabG3C pGC0000 F* | aacaGGTCTCaggctcaacaATGGCTTCTCGGCGGCGAGT | Cloning |
| *AtRabG3C pGB0000 R* | aacaGGTCTCtctgaTTAGCATTCGCACCCAGTTGATCTTTGTTG | Cloning |
| *AtMC2 pGB0000 F* | aacaGGTCTCaaacaATGTTGTTGCTGGTGGACTGCT | Cloning |
| *AtMC2 pGB0000 R* | aacaGGTCTCtagccTAAAGAGAAGGGCTTCTCATATACAG | Cloning |
| *AtMC2$^{C256A}$ F* | TGCCATCGTCGACGCTgcTCATAGTGGTACCGTCATGG | Cloning (Site-directed Mutagenesis) |
| *AtMC2$^{C256A}$ R* | CCATGACGGTACCACTATGAgcAGCGTCGACGATGGCA | Cloning (Site-directed Mutagenesis) |
| *AtSSI4 (AT5G41750) attB1* | GGGGACAAGTTTgtacaaaaaagcaggctCCATGGCTTTGTCTTCTTCTTTGtc | Cloning |
| *AtSSI4 (AT5G41750) attB2* | GGGGACCACTTTGTACAAGaaagctgggtATGAGACTCCATGAGAATTCATC | Cloning |

| | | |
|---|---|---|
| *proSSI4 (At5g41750) F* | TCA AAT CTA AGT CCA CAC AAA GAG | Cloning |
| *proSSI4 (At5g41750) R* | GAG AGA TCA CAA ATG CTT GAA CT | Cloning |
